# Unveiling the role of osteosarcoma-derived secretome in premetastatic lung remodelling

**DOI:** 10.1101/2023.05.06.539690

**Authors:** Sara F.F. Almeida, Liliana Santos, Gabriela Ribeiro, Hugo R.S. Ferreira, Nuno Lima, Rui Caetano, Mónica Abreu, Mónica Zuzarte, Ana Sofia Ribeiro, Artur Paiva, Tânia Martins-Marques, Paulo Teixeira, Rui Almeida, José Manuel Casanova, Henrique Girão, Antero J. Abrunhosa, Célia M. Gomes

## Abstract

Lung metastasis represents the leading cause of osteosarcoma-related death. Progress in preventing lung metastasis is pretty modest due to the inherent complexity of the metastatic process and the lack of suitable models. Herein, we provide mechanistic insights into how osteosarcoma systemically reprograms the lung microenvironment for metastatic outgrowth using metastatic mouse models and a multi-omics approach.

We found that osteosarcoma-bearing mice or those preconditioned with cell-secretome harbour profound lung structural alteration with airways damage, inflammation, neutrophil infiltration, and remodelling of the extracellular matrix with deposition of fibronectin and collagen by stromal activated fibroblasts for tumour cell adhesion. These changes, supported by transcriptomic and histological data, promoted and accelerated the development of lung metastasis. Comparative proteome profiling of the cell secretome and mouse plasma identified a large number of proteins engaged in the extracellular-matrix organization, cell-matrix adhesion, neutrophil degranulation, and cytokine-mediated signalling, which were consistent with the observed lung microenvironmental changes. Moreover, we identified EFEMP1, a secreted extracellular matrix glycoprotein, as a potential risk factor for lung metastasis and a poor prognosis factor in osteosarcoma patients.

## The Paper Explained

### Problem

The high propensity of osteosarcoma to metastasize to the lung is the main reason for treatment failure and cancer-related death. Deciphering the mechanisms driving the metastatic spread is essential to identify potential druggable targets that may halt or prevent metastasis and overcome this adverse scenario of dismal survival.

### Results

We found that metastatic osteosarcoma cells secrete specific biomolecules that travel through the circulation and prepare in advance a unique lung microenvironment for metastasis development. We uncover the ECM remodeling with deposition of cell adhesion promoting proteins by stromal activated fibroblast, and neutrophils recruitment as early pro-metastatic events guiding lung metastasis. Moreover, we identified a secreted extracellular glycoprotein EFEMP1 as a potential prognostic biomarker for lung metastasis in osteosarcoma.

### Impact

This study provides a comprehensive understanding of how primary osteosarcoma systemically reprograms the lung microenvironment for circulating tumor cell colonization and metastasis outgrowth. This knowledge paves the way to establish new strategies to hamper lung metastasis development. Moreover, the newly identified secreted factor EFEMP1 might enhance the identification of osteosarcoma patients with high risk for lung metastasis.

### Synopsis

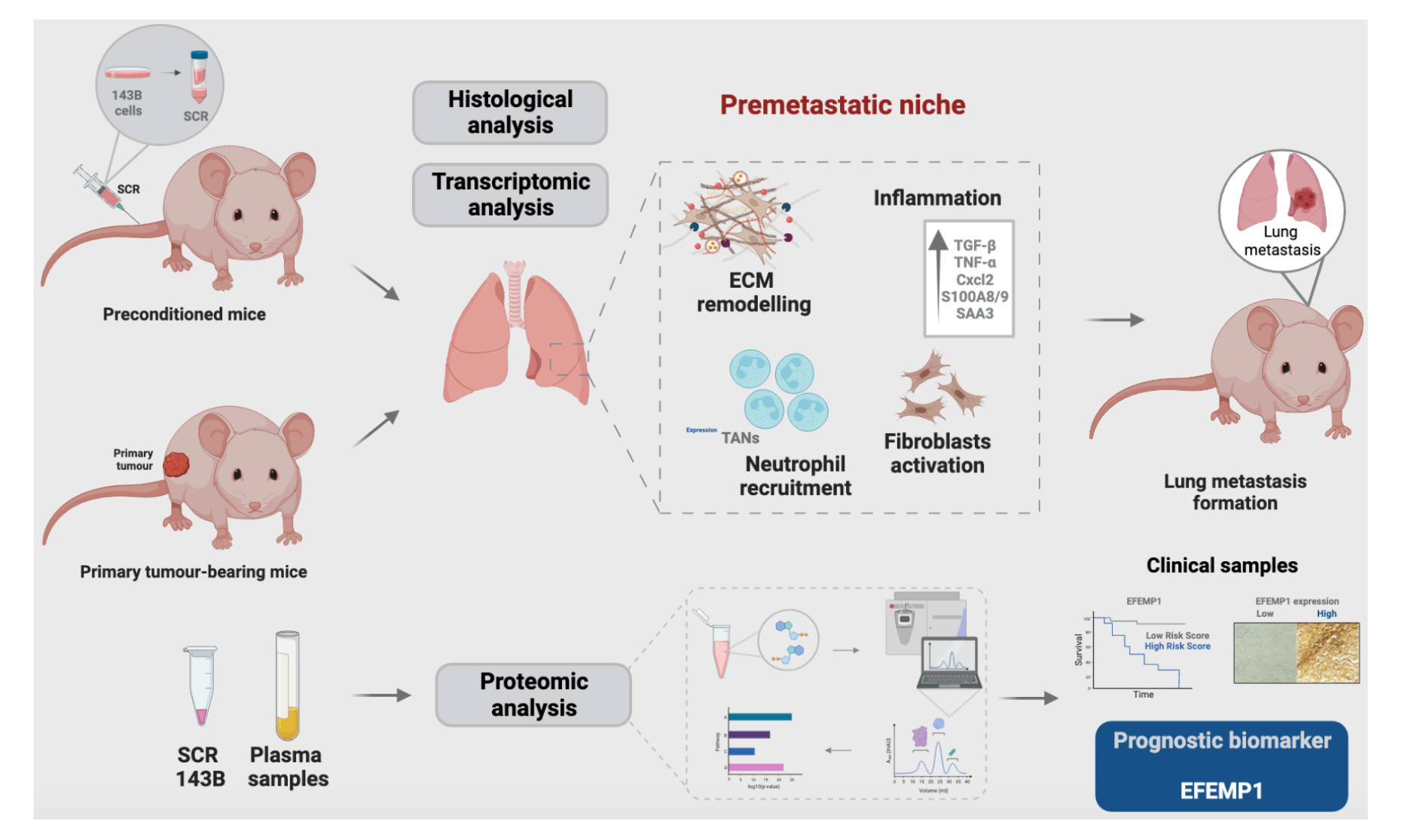

Osteosarcoma-derived secreted factors reprogrammed systemically the lung microenvironment and fostered a growth-permissive niche for the incoming disseminated cells to survive and outgrow into overt metastasis.

- Daily administration of the osteosarcoma cell secretome mimics the secretory dynamics of a growing tumour in mice during the PMN formation;
- Transcriptomic and histological analysis of premetastatic lungs revealed inflammatory-induced stromal fibroblast activation, neutrophil infiltration, and remodelling of the ECM as early-onset pro-metastatic events;
- Proteome profiling identified EFEMP1, an extracellular secreted glycoprotein, as a potential predictive biomarker for lung metastasis and poor prognosis in osteosarcoma patients.
- Osteosarcoma patients with EFEMP1 expressing biopsies have poorer overall survival.

## Introduction

Osteosarcoma (OS) is the most prevalent primary malignant bone tumour that primarily affects children and adolescents (Czarnecka *et al*, 2020; Isakoff *et al*, 2015). OS has a high propensity to metastasize to the lungs, accounting for 90% of metastatic sites, making this the foremost major cause of morbidity and mortality rate (Lindsey *et al*, 2017). The estimated 5-year survival rate of OS patients with localized disease is 70% but drops dramatically to less than 20% in patients who developed metastasis, owing to the ineffectiveness of current therapeutic strategies and late diagnosis (Moukengue *et al*, 2022; Sheng *et al*, 2021).

It is speculated that over 80% of OS patients have undetectable lung micrometastases at initial diagnosis (Lamplot *et al*, 2013) that eventually progress to lethal despite the aggressive multiagent (neo)adjuvant chemotherapy these patients receive. Despite intensive efforts to improve the outcome of OS patients, the survival benefits reported in clinical trials evaluating new therapies are quite marginal as a result of metastatic disease (Harris & Hawkins, 2022). Mechanistically, the development of distant metastasis is a complex and multi-step process distinct from primary tumour formation. The dynamic crosstalk between cancer cells and the local microenvironment is being recognized as a critical regulator of tumour progression and metastasis (Fidler, 2003; Joyce & Pollard, 2009; Neophytou *et al*, 2021). It is now evident that primary tumours prepare in advance and remotely a supportive and receptive microenvironment in specific secondary organs - the so-called pre-metastatic niche (PMN), for the forthcoming tumour cells to adapt and survive (Peinado *et al*, 2017; Sceneay *et al*, 2013). PMN formation encompasses a series of sequential and dynamic events in premetastatic organs primed by tumour-secreted factors and extracellular vesicles (EVs). This process involves a complex interplay of these factors with local stromal cells and tumour-mobilised bone marrow-derived cells (BMDCs) (Liu & Cao, 2016; Wang *et al*, 2021b). Vascular leakage, abnormal extracellular matrix (ECM) remodelling, and immunosuppression have been identified as pro-metastatic events (Liu & Cao, 2016; Peinado *et al*., 2017; Wang *et al*, 2021a).

Therapeutic strategies targeting PMN formation in organ-specific sites offer an opportunity to prevent or suppress metastasis formation and are nowadays a hot topic in cancer research (Li *et al*, 2021; Zhou *et al*, 2020). The identification of potential druggable targets requires a tumour-specific understanding of the cellular and molecular mechanisms involved in establishing organ-specific PMNs. Despite the advances on this topic, sarcomas have been less studied, and the mechanisms enabling the development of organotropic lung metastasis are not clearly understood (Fan *et al*, 2020; Sheng *et al*., 2021).

The cancer secretome is a reservoir of potential biomarkers and signalling biomolecules relevant for tumour progression and metastasis (Stastna & Van Eyk, 2012; Xue *et al*, 2008). This class of proteins has been studied in the search for cancer biomarkers and to understand their mechanistic role in tumour progression (Burns *et al*, 2020; Sirikaew *et al*, 2022). Jerez and colleagues (Jerez *et al*, 2017) conducted a proteomic analysis of the secretome of human OS cell and identified specific proteins in both exosomes and soluble fractions, involved in biological functions (angiogenesis, cellular adhesion, and migration) related to tumour progression and metastasis. Recently, Mazumdar *et al*. (Mazumdar *et al*, 2020a) explored the functional role of OS-derived EVs in driving lung metastatic colonization. Despite having observed a preferential accumulation of EVs in the lungs, the induced stromal changes did not result in an increased tumour burden, suggesting that EVs dictate the organotropism but additional tumour-secreted factors are required to establish a functional PMN (Mazumdar *et al*., 2020a).

In this work, we evaluated how primary OS systemically reprograms the lung microenvironment to establish a permissive pro-metastatic niche for subsequent metastasis formation, using two murine models and a multi-omics approach. Transcriptomic data and tissue analysis revealed ECM remodelling, stromal fibroblast activation, and neutrophil recruitment as the pro-metastatic events that precede lung colonization. Furthermore, we identified EFEMP1, a secreted glycoprotein by tumour cells, as a potential prognostic biomarker for patients with high risk for lung metastasis.

## Results

### Osteosarcoma cell-derived secretome induces transcriptome and tissue structural changes similar to a primary growing tumour

As the metastatic microenvironment plays a pivotal role in facilitating the seeding and outgrowth of disseminated tumour cells, we aimed to examine the lung environmental alterations occurring before lung metastasis formation. In order to establish the role of tumour-derived secreted factors in this process, we used two experimental mouse models. In the first one, animals were implanted subcutaneously with 143B OS cells for primary tumour (PT) formation (hereafter referred to as PT-bearing mice) (Fig 1A). Tumours were allowed to grow until reaching an average volume of 50-60 mm^3^, which took approximately 8-10 days (Fig S1A). To reproduce the continuous secretion of factors by a developing primary tumour, a separate group of animals received daily *i.p.* injections of the secretome (SCR) derived from 143B cells (hereafter referred to as SCR-treated mice) for one week (Fig 1 B). In both conditions, none of the animals showed signs of respiratory distress or weight loss (Fig S2A-B).

**Figure 1.**
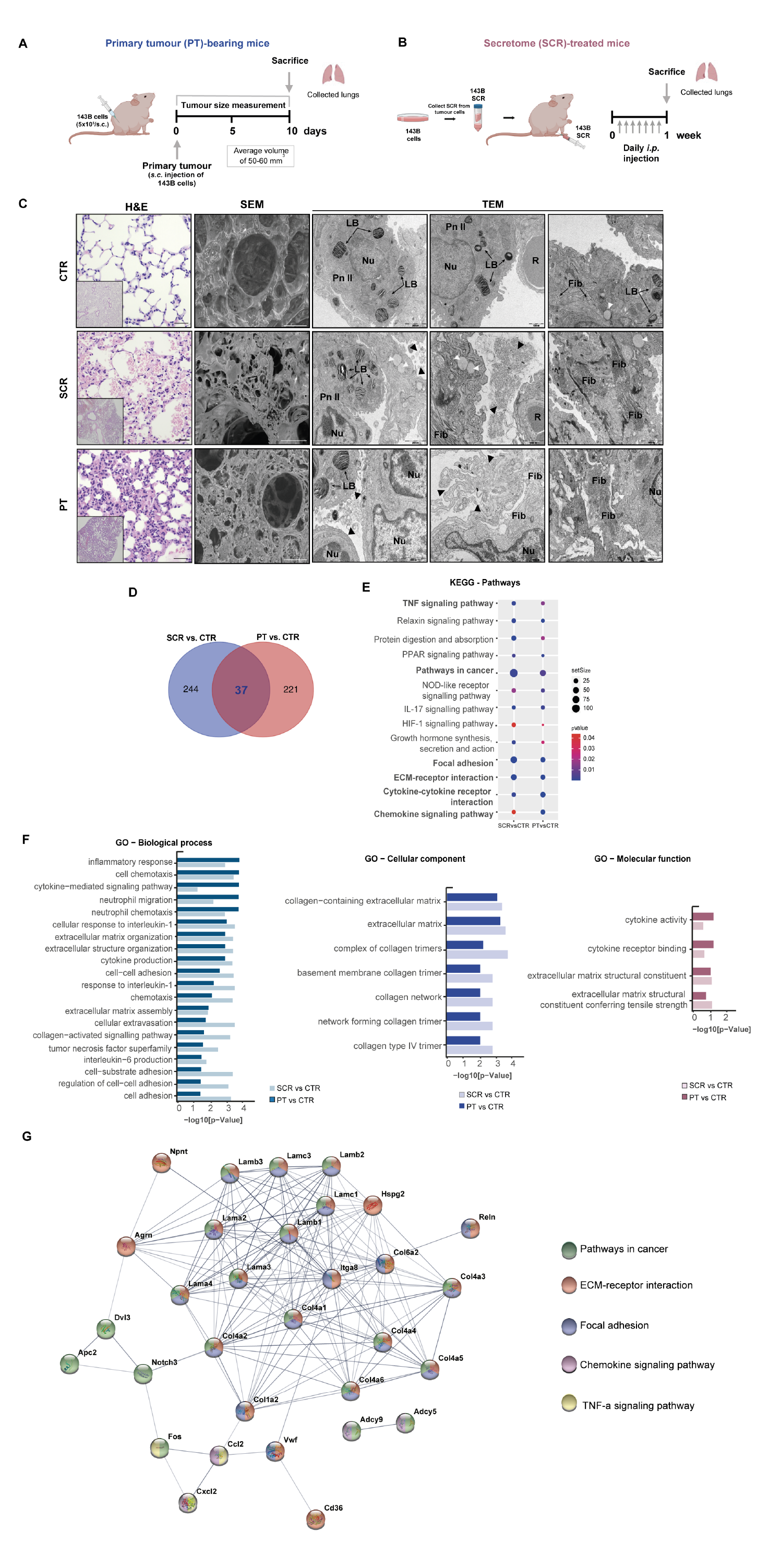
Changes in the transcriptome and lung tissue architecture in mice treated with the secretome or with a primary tumour. A Schematic diagram illustrating the *s.c.* injection of the 143B Luc^+^ cells into the lower flank of mice. Lung tissue was harvested when the primary tumour (PT) reached a maximum volume of 50-60 mm^3^ (PT-bearing mice, n=3-5). B Schematic diagram illustrating the preparation and administration schedule of the143B cell-derived secretome in mice. Animals received daily *i.p.* injections for 1 week (SCR-treated mice, n=3-5). Lung tissue was harvested at the end of the treatment. C Representative images of H&E at x200 magnification (Scale bar: 20 µm), SEM (Scale bar: 40 µm), and TEM (Scale bars: 1000 and 2000 nm) of lung tissue sections from untreated mice (CTR), SCR-treated or PT-bearing mice. Black arrowheads: exudate of protein, vesicles, and fragments of the surfactant; White arrowheads: mucous granules (produced by peribronchial glands); LB, lamellar bodies; Nu, nucleus; R, red blood cells; Pn II: pneumocytes type II (alveolar cells); Fib: fibrosis. D Venn diagram of differentially expressed genes (DEGs) in lungs for each pairwise comparison: SCR vs. CTR and PT vs. CTR. E KEGG pathway enrichment analysis of DEGs. Circle sizes denote the number of genes included in a group and the colour indicates the p-value. F Bar plots depicting the manually curated common Gene Ontology (GO) terms found for the two comparison groups. Biological process (BP), cellular component (CC), and molecular function (MF) of altered genes reporting the intersections in lungs from SCR vs. CTR and PT vs. CTR. G Representative protein-protein interaction (PPI) network, constructed with the common DEGs, using the STRING database.

The lungs were subsequently harvested for histological and electron microscope examinations. Histopathological analysis of formalin-fixed paraffin-embedded (FFPE) lung tissue from PT-bearing and SCR-treated mice demonstrated severe damage to the alveolar structure, with significant septum thickening compared to vehicle-treated control (CTR: 7.92 ± 0.40 µm; SCR: 15.79 ± 1.13 µm; PT: 24.25 ± 2.01 µm, p<0.001), reduced airspace areas, and inflammatory cell infiltration, in comparison to the preserved lung parenchyma from control mice (Fig 1C). No microscopically detectable lung metastasis was found in the lungs of PT-bearing mice at the time of the endpoint.

Scanning electron microscopy (SEM) images revealed marked changes in the overall alveolar architecture with condensation of the connective tissue framework surrounding the alveolar spaces in the lungs of both PT-bearing and SCR-treated mice. Furthermore, transmission electron microscopy (TEM) images confirmed these observations and showed a disorganized and severely damaged pulmonary structure with the leakage of proteins, vesicles and surfactant (arrowheads) into the alveolar space. Additionally, septum thickening, nucleus with altered morphology and fibrosis, i.e. overproduction and accumulation of the ECM proteins, were evident in comparison to healthy controls.

These findings revealed early-onset structural changes in the lung parenchyma, likely induced by the continuous release of distant tumour-secreted factors. Furthermore, the effects resulting from the daily administration of SCR closely resemble those observed in the host with the tumour, indicating a systemic tumour-mediated effect.

Aiming at understanding the transcriptome alterations underlying the lung histological and architectural changes, we performed a comparative RNA sequencing-based transcriptome profiling of the lung tissue collected from healthy, SCR-treated, or PT-carrying mice.

Among the 16,138 transcripts, we found a total of 502 differently expressed genes (DEGs) between controls and the experimental groups, using a cut-off of p-value<0.05 and log fold change (FC)>|1|. From these, 37 were shared between the two compared groups as shown in the Venn diagram (Fig 1D). Gene set enrichment analysis (GSEA) identified enrichment of pathways in cancer, ECM-receptor interaction, focal adhesion, chemokine signalling pathway, and cytokine-cytokine receptor interactions (Fig 1E). Gene ontology (GO) functional enrichment analysis of the common DEGs confirmed that they significantly contributed to the enrichment in several biological processes (BP) terms including inflammatory response, cell chemotaxis, cytokine-mediated signalling pathway, neutrophil migration and chemotaxis, cellular response to interleukin-1, ECM and structure organization, cytokine production, cell-cell adhesion, cellular extravasation, interleukin-6 production and cell adhesion (Fig 1F). Common DEGs were also enriched in cellular components (CC) terms including the ECM and collagen and in molecular functions (MF) terms comprising the ECM structural constituents, cytokine receptor binding and cytokine activity (Fig 1F). Furthermore, a protein-protein interaction (PPI) network analysis of the overlapping 37 DEGs identified 159 edges among 33 nodes (PPI enrichment p-value<1.0×10^-16^, using the String platform), with genes such as collagens (Col1, Col4 and Col6), laminins (Lama, Lamb and Lamc) and integrin alpha-8 as a part of the main cluster of ECM-receptor interactions, focal adhesion and pathways in cancer (Fig 1G). Collagen, proteoglycans (such as versican and hyaluronan) and glycoproteins (such as laminins, elastin and fibronectin) are the core matrisome proteins of the ECM maintaining the complex tissue architecture (Winkler *et al*, 2020). Changes in ECM remodelling have implications on cellular signalling networks since ECM components can function as pro-inflammatory stimuli and serve as ligands for various molecules and cell surface receptors, such as integrins, likely impacting metastasis formation (Bonnans *et al*, 2014; Popova & Jücker, 2022; Winkler *et al*., 2020). The transcriptomic data strongly support this hypothesis since several ECM-related pathways were significantly changed by the growing tumour and secretome.

### Osteosarcoma-secreted factors cause changes in the lung ECM and the immune and inflammatory landscape during PMN formation

After identifying the core biological pathways through GSEA, we performed a heatmap-based analysis of the relative expression of transcripts linked to ECM remodelling, inflammation, and immune cell recruitment-related pathways between the control and experimental groups. Transcripts with significant expression changes (p-value < 0.05) compared to control mice are marked with a black border (Fig 2A-B).

**Figure 2.**
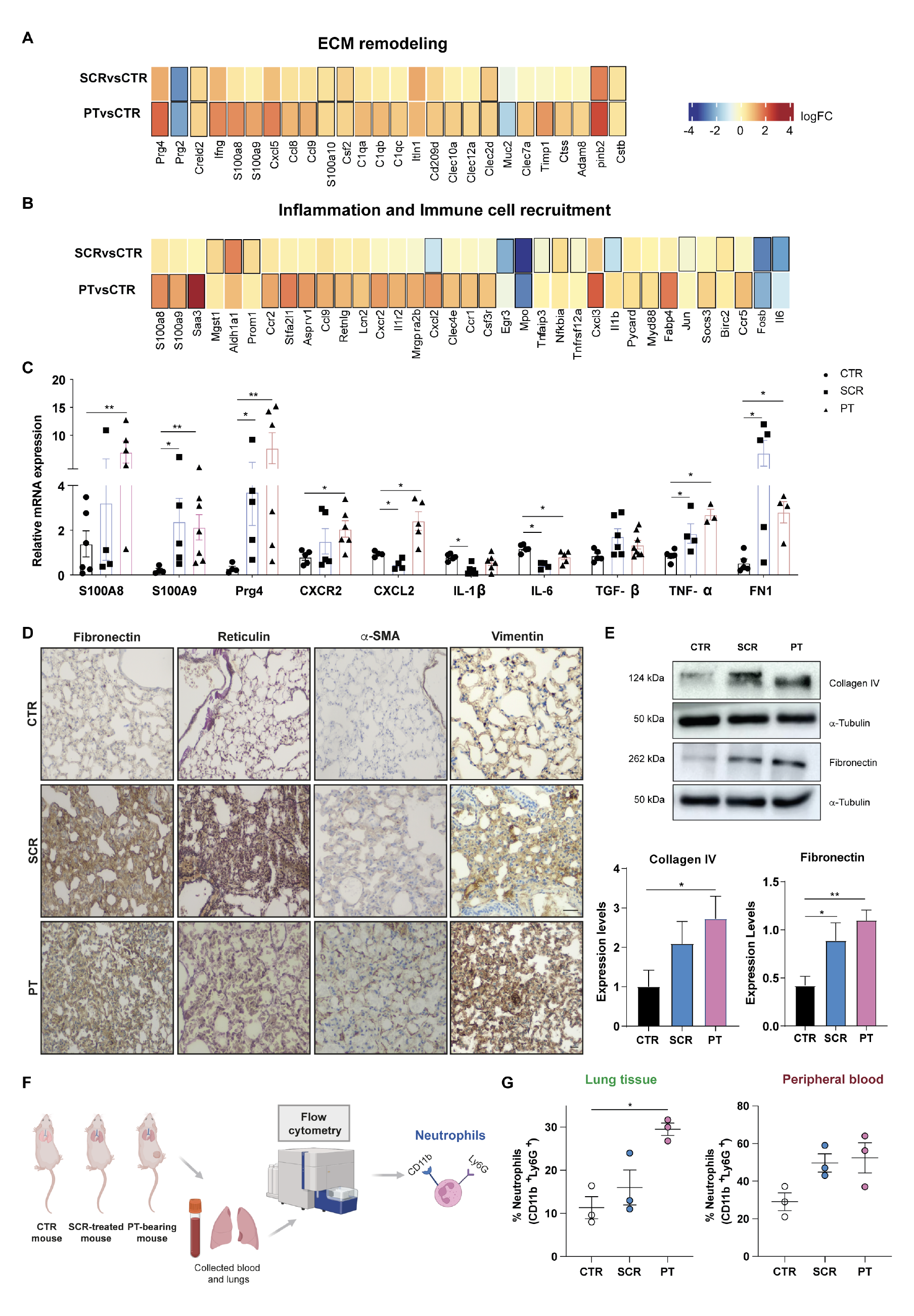
Lung microenvironmental changes in response to osteosarcoma-secreted factors. A, B Heatmaps of transcripts encoding genes involved in ECM remodelling, inflammation, and immune cell recruitment in lungs from SCR-treated or PT-bearing mice compared with controls. C qRT-PCR analysis of S100A8/A9, Prg4, Cxcr2, Cxcl2, IL-1β, IL6, TGF-β, TNF-α, and FN1 genes in lungs from SCR-treated mice or PT-bearing mice compared with controls. (n=3-7, per group). D Representative images of fibronectin, reticulin, α-SMA, and vimentin immunostaining at x200 magnification (Scale bar: 30 µm) in lung sections from CTR mice, SCR-treated mice, or carrying a PT E Western blot analysis of fibronectin and collagen type IV. Expression levels with graphic quantification. F Schematic diagram illustrating the analysis of lung-infiltrating neutrophils and the peripheral blood in mouse models by flow cytometry. G Flow cytometric quantification of infiltrating neutrophils in the lungs and in the peripheral blood of control, SCR-treated and PT-bearing mice (n=3, per group). Data information: Data are presented as mean ± SEM from 3-7 independent biological samples. *p<0.05, **p<0.01 compared to control lungs (Mann-Whitney test (C, E and G)).

Lung transcriptomic analysis revealed a higher number of significantly dysregulated genes in mice with PT compared to those treated with the SCR, although the overall trend is similar (Fig 2A-B). Secretome was collected from 143B cells in a two-dimensional monoculture and, despite being administered daily, it may not entirely reproduce the secretion rate of a growing tumour, nor the influence of its surrounding stromal microenvironment. This could explain for the relatively less pronounced effects observed in the SCR-treated mice.

The heatmap of ECM-related genes displayed 26 deregulated genes (Fig 2A) including the upregulation of the proteoglycan 4 (Prg4) with an anti-inflammatory role (Alquraini *et al*, 2015; Iqbal *et al*, 2016), the downregulation of the proteoglycan 2 (Prg2) known to exert tumour-suppressor functions (Carpino *et al*, 2019) and the upregulation of the cysteine-rich with EGF-like domains protein 2 (Creld2) that have been linked to tumour progression via fibroblast reprogramming (Boyle *et al*, 2020). Amongst ECM-related genes, several ECM-affiliated, secreted factors and ECM regulators were found dysregulated, such as the C1q complement components of collagen-like structures and C-type lectin-like receptors, known to act as tumour-promoting factors and inflammatory drivers (Bulla *et al*, 2016; Chiffoleau, 2018). The S100 calcium-binding proteins A8/9 (S100A8/A9), the chemokines Cxcl5, Ccl8 and 9, and the colony-stimulating factor (Csf) 2 in the category of ECM-secreted factors, are involved in cell proliferation, invasion and growth, inflammation and function as chemo-attractants for the recruitment and activation of myeloid cells (Chen *et al*, 2014; Marcuzzi *et al*, 2018; Shi *et al*, 2006). Indeed, S100A8 and A9 were found to regulate the expression of serum amyloid A (Saa) 3, which acts as a positive-feedback modulator for chemoattractant secretion for CD11b^+^ myeloid cells (Hiratsuka *et al*, 2008). The ECM regulators plasminogen activator inhibitor type 2 (Serpinb2) and the cystatins (Cst) b/s, also upregulated, have been recognized as reliable markers for predicting tumorigenicity (Lee *et al*, 2019; Wang *et al*, 2014).

The heatmap of genes with inflammation and immune cell recruitment revealed 35 deregulated genes (Fig 2B), from which we highlighted chemokines (Ccl9, Cxcl2 and Cxcl3), chemokine receptors (Ccr2, Cxcr2 and Ccr1) and the SAA and S100A8/A9, that cooperate in the recruitment of specific myeloid populations including neutrophils, monocytes and macrophages. The proteolytic enzyme myeloperoxidase (Mpo), considered a marker of neutrophil degranulation (Aratani, 2018), is downregulated together with the early growth response 3 (Egr3), a suppressor gene for tumour initiation and progression (Zhang *et al*, 2017). The upregulated transcripts microsomal glutathione S-transferase (Mgst1) and the aldehyde dehydrogenase 1 (Aldh1a1) along with lipocalin-2 (Lcn2), have been linked to the immunosuppressive function of myeloid-derived suppressor cells (MDSCs) to facilitate tumour progression (Kaplan *et al*, 2006; Liu *et al*, 2021; Sokol & Luster, 2015; Yan *et al*, 2022) as the fatty acid-binding protein 4 (Fabp4) recently associated with metastatic aggressiveness (Marcuzzi *et al*., 2018; Zhang *et al*, 2019).

Several of the differentially expressed transcripts were confirmed by qRT-PCR in lung tissue lysates, including S100A8/A9, Prg4, Cxcr2, Cxcl2, IL-1β, and IL6, and the results were consistent with RNA-seq data (Fig 2C). Additionally, we examined the mRNA expression of TGF-β, TNF-α, and fibronectin, known for their immunosuppressive, pro-angiogenic, and ECM regulatory roles, respectively, which are also upregulated in the lungs of PT-bearing and SCR-treated mice, compared to the controls.

Immunohistochemical analysis of lung tissue sections revealed a marked increase in the deposition of two major ECM components, fibronectin and reticulin fibers, as well as upregulation of the activated fibroblast marker α-SMA and vimentin in both SCR-treated and PT-bearing mice, compared to the control group (Fig 2D). It was also observed an increase in type IV collagen, (the major component of the basement membrane) by western blot, as well as of fibronectin (Fig 2E). These findings support the transcriptomic evidence of lung ECM remodelling during PMN formation.

The transcriptome profile also identified biological processes and differentially expressed genes that cooperate in the recruitment of myeloid cells. Considering that, we conducted a flow cytometry analysis of infiltrating neutrophils in lung tissue and in the peripheral blood of untreated, SCR-treated or PT-bearing mice (Fig 2F). The results showed a significant increase in the percentage of infiltrating neutrophils (CD11b^+^ Ly6G^+^) in the lungs of PT-bearing mice relative to age-matched untreated mice (Fig 2G). The same trend was observed in the lungs and peripheral blood of SCR-treated mice, although without statistical significance compared to healthy controls. Neutrophils are the most abundant leukocytes in circulation and have been recognized as part of the immune response to facilitate metastatic spread (Wu *et al*, 2020). Depending on the microenvironment, neutrophils can have an anti-tumorigenic N1 or a pro-tumorigenic N2 phenotype (Masucci *et al*, 2019). We were unable to figure out the real phenotype of the lung-infiltrating neutrophils, but the increased mRNA levels of TGF-β, one of the main modulators of neutrophil polarization, suggest an N2-like phenotype. Moreover, the transcriptomic analysis revealed an upregulation of Lcn2, a secreted glycoprotein produced by N2-neutrophils and implicated in promoting metastasis (Ören *et al*, 2016; Tyagi *et al*, 2021) alongside a marked downregulation of Mpo in the premetastatic lungs, further supporting this hypothesis.

Altogether, these findings suggest that tumour-secreted factors instigate inflammatory signalling pathways in the lung microenvironment with concurrent ECM remodelling and neutrophil infiltration, as part of the PMN formation.

### Activated lung fibroblasts in premetastatic lungs drive fibronectin and collagen deposition and favour the adhesion of OS cells to the lung ECM

As fibroblasts constitute the most abundant cells in the lung interstitium and are the major producers of ECM components, we hypothesized that tumour-secreted factors can induce ECM remodelling through alterations in the phenotype and function of lung fibroblasts. To address that, we conducted a phenotypic profiling analysis of lung fibroblasts isolated directly from fresh lungs of SCR-treated and PT-bearing mice as schemed in Fig 3A. Fibroblasts from healthy age-matched mice were used as control and designated as normal fibroblasts (NFs). Immunostaining showed an upregulation of α-SMA and FAP, two classical markers of activated fibroblasts, as well as of the intermediate filament vimentin, in fibroblasts from SCR-treated animals or carrying a PT, and were hereafter referred to as normal activated fibroblasts, NAFs^SCR^ or NAFs^PT^, respectively. The strong immunoreactivity for fibronectin in NAFs^SCR^ and NAFs^PT^, further supports the contribution of stromal fibroblasts in lung ECM remodelling, as previously described. In addition, staining with phalloidin, a marker of F-actin filaments, showed changes in actin organization, characterized by an irregular and disorganized shape and shortening of filament structures (Fig 3B) in fibroblasts derived from PT-bearing and SCR-treated animals. These phenotypic changes were accompanied by an increase in cell motility, as assessed by a wound healing assay (Fig S3A-B).

**Figure 3.**
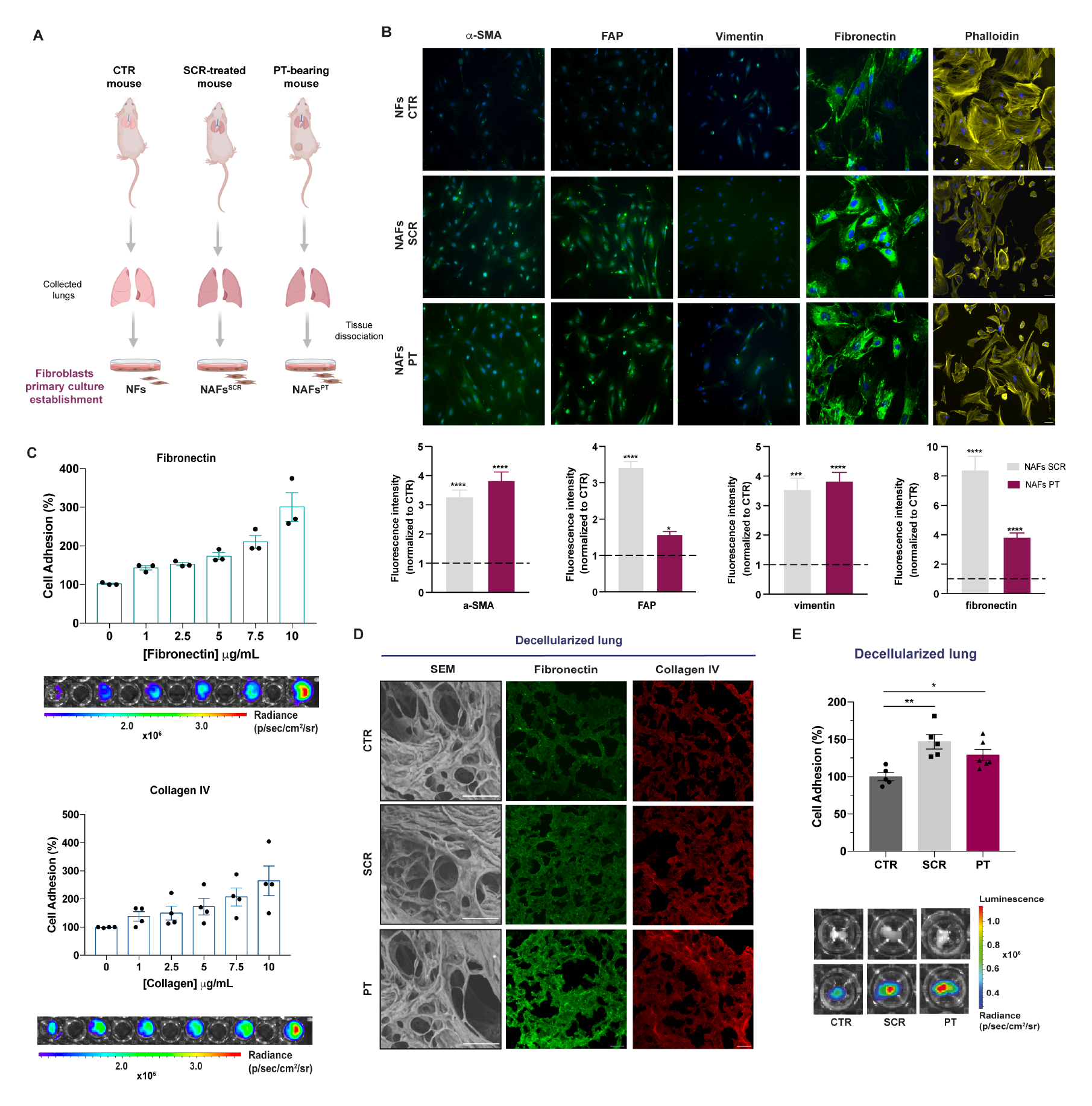
Increased deposition of fibronectin and collagen by activated lung fibroblasts favour the adhesion of OS cells to the lung ECM. A Schematic diagram of the establishment of the primary cultures of fibroblasts isolated from the lungs of untreated (NFs), SCR-treated (NAFs^SCR^) and PT-bearing mice (NAFs^PT^). B Representative image of immunofluorescence staining for α-SMA, FAP, vimentin, fibronectin and phalloidin at x20 magnification (Scale bar: 50 µm) in primary cultures of fibroblasts from CTR, SCR-treated or PT-bearing mice. Immunofluorescence quantification of α-SMA, FAP, vimentin and fibronectin (n=3, per group). C Relative adhesion of 143B cells to increasing concentrations of fibronectin and collagen type IV ranging from 1 to 10 µg/mL (n=3-4, performed in triplicate). Representative bioluminescence images of cell adhesion (143B cells) to different concentrations of fibronectin and collagen IV. D Representative SEM images (Scale bar: 50 µm) of decellularized lung sections and fibronectin and collagen immunostaining at x20 magnification (Scale bar: 50 µm) in decellularized lung sections from CTR mice, SCR-treated mice, or carrying a PT. E Relative adhesion of 143B cells to decellularized lung sections from CTR, SCR-treated, or PT-bearing mice (n=5). Representative pictures of the decellularized fragments in the wells (upper row) and bioluminescent images of adhered cells (lower row). The bioluminescent signal is represented as radiance (p/s/cm^2^/sr). Data Information: Data are presented as mean±SEM. *p<0.05 and ****p<0.0001 are significantly different when compared to NFs (Kruskal-Wallis (B)); *p<0.05 and **p<0.01 when compared to healthy decellularized lungs (One-way ANOVA (E)).

Given that fibronectin is a cell-adhesive glycoprotein known to favour the adhesion of tumour cells, we hypothesized that the secretome-induced upregulation of fibronectin could promote further lung colonization by OS cells. To address that, we performed a cell adhesion assay of 143B cells on fibronectin-coated plates using coating concentrations ranging from 1 to 10 μg/mL. The results showed a gradual rise in the percentage of adhering cells with increasing concentrations of fibronectin. Similar results were observed on collagen IV-coated plates, demonstrating that these proteins act as substrates for cell attachment (Fig 3C).

Encouraged by these findings, we next set out to evaluate the adhesion of 143B cells to decellularized lung fragments from SCR-treated or PT-bearing mice. The immunofluorescence staining confirmed the increased deposition of fibronectin and collagen IV in the decellularized fragments of animals treated with the SCR or with a PT (Fig 3D), in line with IHC and WB data (Fig 2D, E). The 143B cells were seeded in plates coated with the decellularized lung scaffolds and allowed to attach for 10 min. Our results revealed a significant increase in the percentage of 143B cells attached to the lung scaffolds of mice treated with SCR (p<0.01) and PT-bearing mice (p<0.05), with an increase of approximately 50% compared to control animals (Fig 3E). Overall, our data demonstrate that factors secreted by OS cells induce remodelling of the ECM lung, facilitates the adhesion of cancer cells, likely preceding metastasis formation.

Furthermore, the immunostaining of decellularized fibroblast cell sheets revealed an increase in the staining intensity for fibronectin and collagen IV (Fig S4A), which confirms the contribution of activated fibroblasts in ECM remodelling and cell adhesion.

### Tumour secretome-induced microenvironmental changes in pre-metastatic lungs promote and accelerate lung metastasis formation

Up to this point, our data indicates that the secretome of OS cells induces ECM remodelling and immune cell infiltration in the lungs, fostering PMN formation. To further investigate whether OS secretome promotes and/or accelerates the formation of lung metastasis *in vivo*, we resorted to an experimental lung metastasis mice model. For that, preconditioning of the animals by daily administration of the 143B secretome was performed for one week, followed by the *i.v.* injection of tumour cells into the tail vein (Fig 4A). Non-preconditioned mice were used as controls (Fig 4B). Lung metastasis formation was monitored weekly by bioluminescence imaging (BLI) for over 60 days.

**Figure 4.**
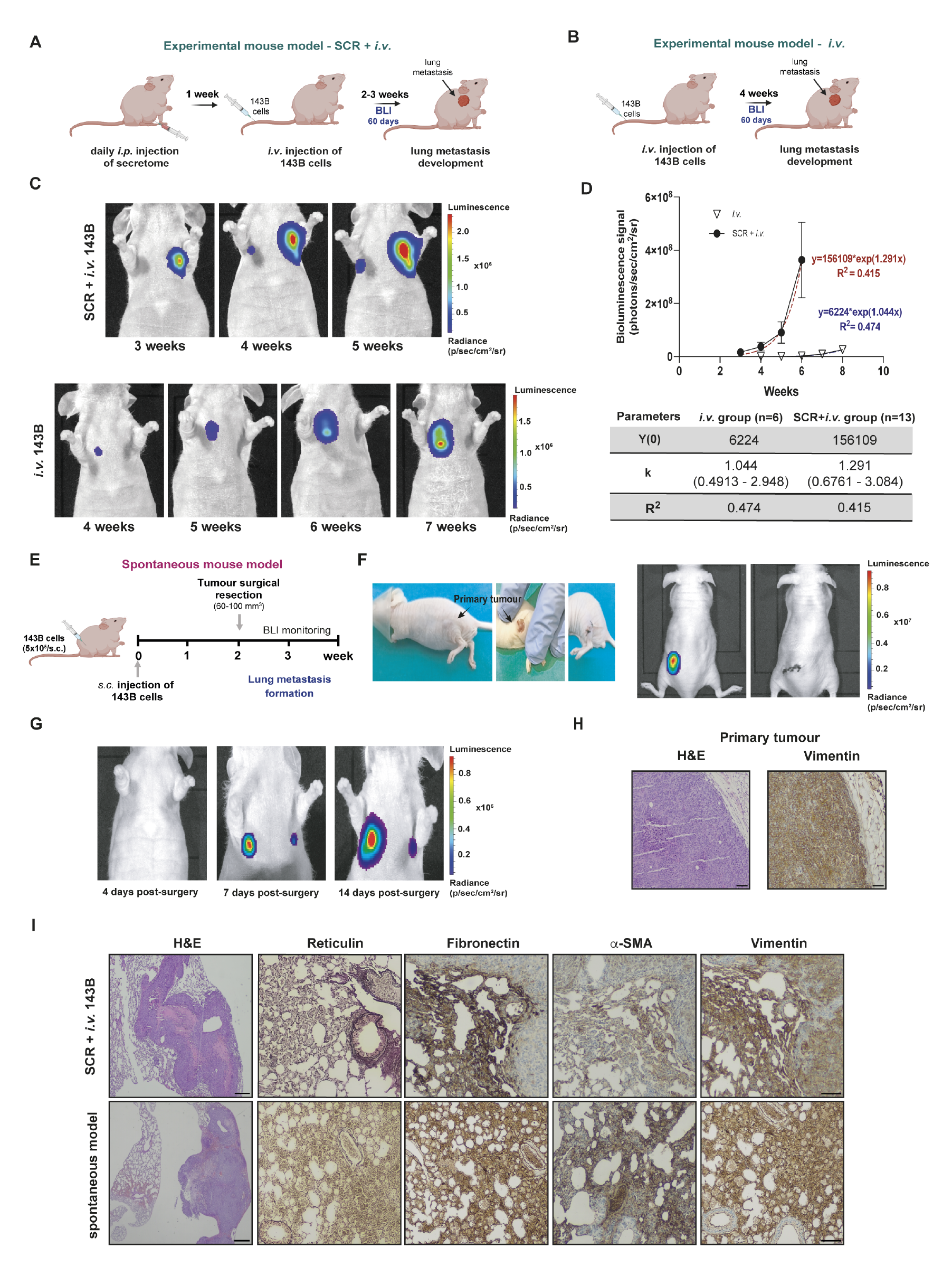
Osteosarcoma-induced PMN formation promotes and accelerates the formation of lung metastasis. A Schematic diagram of the experimental model of lung metastasis. Mice were treated with 143B cells-derived secretome (SCR) for 1 week, followed by *i.v.* administration of 143B Luc^+^ cells into the tail vein (SCR+*i.v.* group). B Schematic diagram of the experimental model of lung metastasis. Mice received only the *i.v.* injection of 143B Luc+ cells without pre-treatment with the SCR (*i.v.* group). C Representative bioluminescence images of lung metastasis formation in pre-treated (SCR+*i.v. group)* and untreated (*i.v.* group) mice with secretome before the *i.v.* injection of 143B cells. D Exponential fitting of the bioluminescence signals (photons/second) of metastatic lesions over time in the *i.v.* group (ς n=6) and the SCR+*i.v.* group (• n=13 mice), and corresponding kinetic parameters. E Schematic diagram of the spontaneous metastatic mouse model. Animals were injected subcutaneously with the 143B cells. After reaching a volume of 60-100 mm^3^, the tumour was excised, and the animals were monitored by BLI for lung metastasis formation. F Images of the surgical resection of a primary tumour with a volume of 60 mm^3^ and bioluminescence images before and after the excision of the tumour. G Representative bioluminescence images at 4, 7 and 14 days after surgical resection of the primary tumour. H Histological H&E images at x100 magnification (Scale bar: 30 µm) and immunostaining for vimentin at x100 magnification (Scale bar: 30 µm) of the resected tumour. I Histological H&E images at x40 magnification (Scale bar: 20 µm) and IHC staining for reticulin, fibronectin, α-SMA, and vimentin at x100 magnification (Scale bar: 30 µm) of lung metastatic lesions in both experimental and spontaneous mouse models. Data Information: Data are presented as mean ± SEM.

All preconditioned animals developed lung metastasis, detected in the second-third weeks after cells inoculation, that rapidly progressed to larger and multilobular lesions. In contrast, only 55% of the untreated group, developed metastasis by the fourth week, mostly unilobular, and with a slower growth kinetic, never reaching the size of the treated group (Fig 4C). The parameters estimated from the exponential fitting of the bioluminescent signals confirmed the faster growth rate of metastatic lesions in animals preconditioned with the secretome of OS cells (Fig 4D). The development of metastasis in some untreated animals was probably due to the highly aggressive nature of 143B cells. Nonetheless, the metastatic rate was substantially lower, and the lesions were smaller and less invasive compared to the preconditioned group.

We also established a spontaneous metastatic animal model to assess whether the PMN formation induced by an engrafted primary tumour dictate the innate propensity of OS to metastasize to the lung. A primary tumour was induced by subcutaneous injection of 143B cells into the lower flank and allowed to grow until it reached a maximum volume of 60-100 mm^3^ (Fig 4E). The tumour was then surgically excised, and the animals were monitored for lung metastasis formation. BLI images after surgery confirmed the complete removal of the primary tumour (Fig 4F) and the absence of lung metastasis or metastasis in any other organs. Four days post-surgery, there were no apparent signs of lung metastasis, but a week later, two nodules were detected, which progressively evolved over the following two weeks (Fig 4G). These results suggest that tumour cells had already spread throughout the circulation at the time of surgery, which were able to colonize the permissive lung microenvironment induced by the primary tumour, allowing for their survival and metastatic outgrowth. The spontaneous formation of distant lung metastasis further confirms the organotropism of OS for this organ, which is believed to be attained by secreted factors by the engrafted OS cells.

The histopathological examination confirmed the high-grade malignancy of both primary OS (Fig 4H) and the lung metastasis in both experimental and spontaneous models (Fig 4I), as evidenced by the marked nuclear pleomorphism, the high mitotic rate and intralesional necrotic areas The silver staining of lung sections revealed an increased deposition and rearrangement of fibrillar collagens surrounding the metastatic lesions and around the bronchi and blood vessels, consistent with a fibrotic response. Additionally, IHC showed strong immunoreactivity for fibronectin and α-SMA in the stromal regions in the periphery of the neoplastic lesions, which reveals the presence of activated fibroblasts in the lung. The mesenchymal marker vimentin, also known as a fibroblast-intermediate filament, exhibited strong immunoreactivity in stromal and metastatic cells confirming the mesenchymal origin of the latter.

Overall, the findings confirmed that lung alterations induced by the secretome or grafted OS cells create a favourable environment for the survival and outgrowth of metastatic cells and undergo continuous remodelling.

### Analysis of the proteomic profile of cell secretome and mouse plasma identified EFEMP1 as a potential prognostic biomarker in osteosarcoma

Since the effects of the daily administration of the SCR on a premetastatic lung were quite similar to those elicited by a local primary tumour, we performed a comparative mass spectrometry-based label-free quantitative proteomic analysis of the 143B secretome and the plasma from PT-bearing mice to identify potential common mediators predictive of lung metastasis formation. The secretome collected from a non-metastatic OS cell line (MG-63) (Ren *et al*, 2015) and the plasma from healthy mice were used as controls. We confirmed the non-metastatic potential of MG-63 cells in mice, as none of the animals *i.v.* injected with these cells developed metastases (Fig S5A).

A list of differentially expressed proteins (DEPs) of 143B *versus* MG-63 OS cells and plasma from healthy mice *versus* PT-bearing mice was generated, using an FDR-adjusted p-value ≤ 0.05.

A total of 2,595 proteins were identified in the secretome of OS cells with 109 being differentially expressed between the 143B and MG-63. In the mouse plasma, we focused on the proteins of human origin, since it represents what has been secreted by human OS-induced tumours. Out of these, we identified 195 differentially expressed between the plasma samples of PT-bearing mice and those of healthy controls.

A GO enrichment analysis was performed on the DEPs between the two pairwise comparisons to identify shared biological processes and molecular functions. The largest number of identified proteins were particularly enriched in neutrophil degranulation, platelet degranulation, ECM organization, and cell adhesion (as depicted in Fig 5A-B).

**Figure 5.**
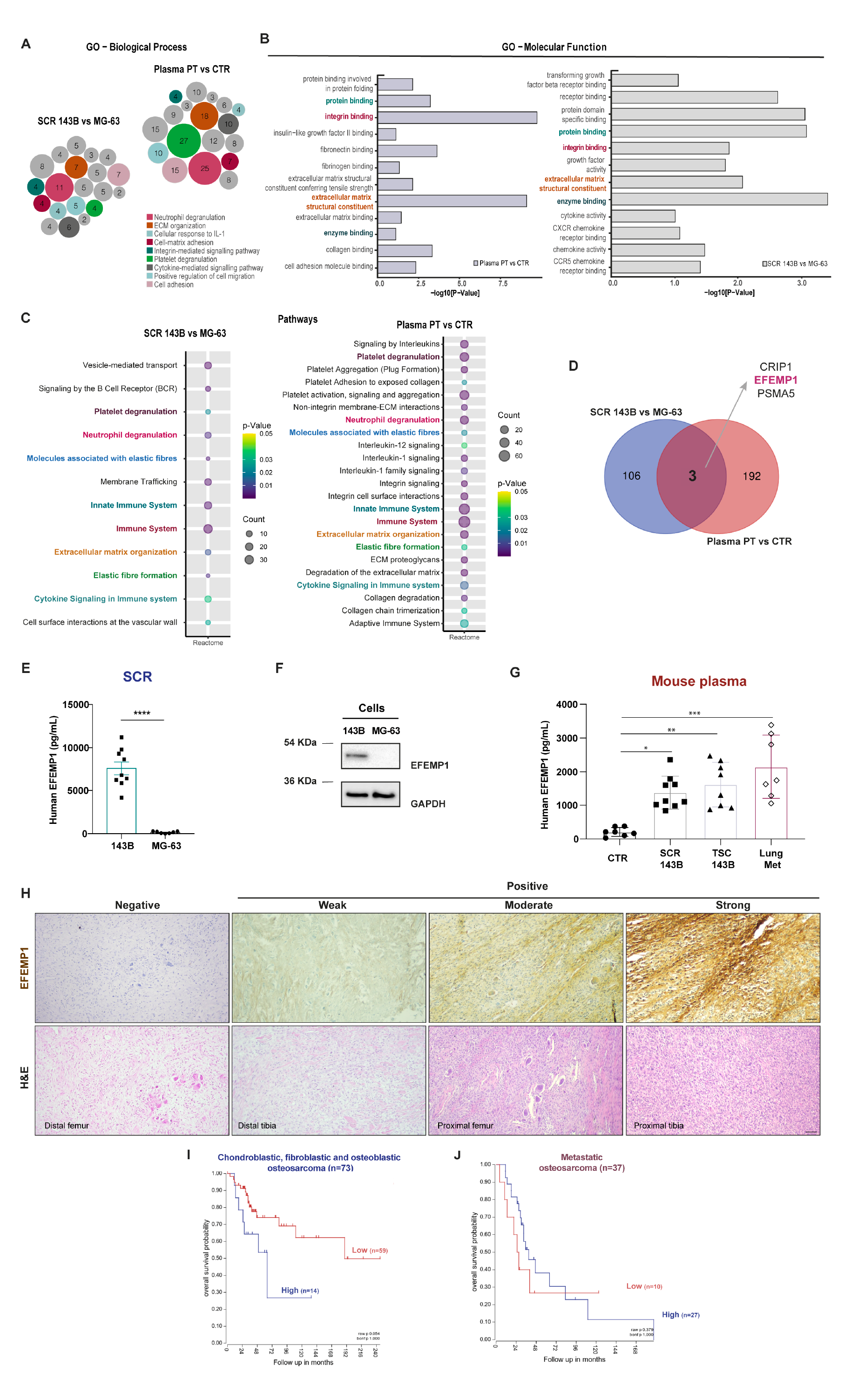
Proteomic analysis identified EFEMP1 as a potential metastatic-related biomarker in osteosarcoma. A-B Gene ontology analysis (GO) output. Biological process (BP) and molecular function (MF) of differently expressed proteins (DEPs) in the two pairwise-comparison groups (SCR 143B vs MG-63; Plasma PT vs. CTR). Circle sizes in BP denote the number of genes involved in the process. C Reactome pathway enrichment analysis of identified proteins in the two pairwise-comparison groups. Circle sizes denote the number of genes included in a group and colour indicates the p-value. The common pathways are highlighted. D Venn diagram showing specific and common proteins among the two pairwise groups: SCR 143B vs. MG-63 and Plasma PT vs. CTR. E EFEMP1 levels in the secretome of the metastatic 143B and non-metastatic MG-63 OS cells. F Representative western blot of EFEMP1 in the metastatic 143B and non-metastatic MG-63 OS cells. G Plasma levels of EFEMP1 in control mice (CTR), mice treated with the 143B secretome (SCR), bearing a primary tumour (PT) or with lung metastasis (Lung Met). H Representative images of EFEMP1 staining and H&E in biopsy samples of high-grade OS patients at x100 magnification (Scale bar: 50 μm). I, J Kaplan-Meier analysis of overall survival in chondroblastic, fibroblastic and osteoblastic OS patient samples (n=73 patients) and with metastatic disease (n=37 patients). Scan cut-off was used to group samples into high (blue) and low (red) EFEMP1 expressions. p-values were determined by a log-rank test. Data Information: Data are presented as mean±SEM, from 7-8 independent experiments. **** p<0.0001 were significantly different when compared with the SCR from MG-63 (unpaired t-test (E)); *p <0.05, **p <0.01 and ***p <0.001 were significantly different when compared with the values present in the plasma from healthy mice (Kruskal-Wallis test (G)).

The Reactome enrichment analysis identified 8 common overlapping pathways engaged in platelets degranulation, neutrophil degranulation, molecules associated with elastic fibres, innate immune system, immune system, ECM organization, elastic fibre formation and cytokine signalling in the immune system (Fig 5C). All of these pathways are known to be involved in the PMN formation (Jablonska *et al*, 2017; Li *et al*, 2020; Mirzapour *et al*, 2023; Yuan *et al*, 2023).

The Venn diagram in Figure 5D, identified three overlapping DEPs between the two pairwise comparisons, which include the epidermal growth factor-containing fibulin-like extracellular matrix protein 1 (EFEMP1, also called fibulin-3), cysteine-rich protein 1 (CRIP1) and proteasome subunit alpha type-5 (PSMA 5). Among these proteins, EFEMP1 was found to be the most abundant in both the secretome and in the mouse plasma, exhibiting consistently high-intensity peak values and greater protein coverage across all replicates. EFEMP1 is a glycoprotein broadly expressed in various tissues during development and adulthood. As a crucial component of basement membranes, EFEMP1 plays a crucial role in preserving the structural integrity and stability of the ECM (Hu *et al*, 2019). Considering the observed alterations in the ECM components in the lungs of mice challenged with the secretome or carrying a PT, we reasoned that secreted EFEMP1 could play an important role in this process. To address that, we started by validation of our proteomic approach by measuring the protein expression and secreted levels of EFEMP1 by OS cells *in vitro*.

The ELISA and western blot analysis provided confirmation of the exclusive expression and secretion of EFEMP1 by 143B cells, with negligible levels observed in the non-metastatic MG-63 cell line, thereby supporting the proteomic data (Fig 5E, F). Moreover, EFEMP1 was detected in the plasma of animals that were treated with the secretome or carrying a PT, as well as in those with established lung metastasis, with higher levels observed in the latter (Fig 5G). The use of a human EFEMP1 ELISA kit points to the 143B cells as the source of the plasmatic EFEMP1 levels. Importantly, this protein is enrolled in 3 out of the 10 common biological processes, specifically ECM organization, molecules associated with elastic fibres and elastic fibre formation, suggesting it may contribute to ECM remodelling during metastasis.

To explore the potential prognostic significance of EFEMP1 expression in OS patients, we conducted a retrospective study on a small cohort of 29 patients without evidence of metastasis at the time of diagnosis. Archived paraffin-embedded biopsy specimens of the primary tumour were immunohistochemically stained for EFEMP1 to investigate its association with the occurrence of lung metastasis during the follow-up period. Of the 29 patients, 12 (41.4%) experienced lung metastasis, and 50% of those patients had positive biopsy specimens for EFEMP1 with a disease-free interval (DFI) of 38.5 months (range: 6 – 77 months). The other 6 patients who tested negative for EFEMP1 had a longer DFI of 44 months (range: 15 −159 months). Among the 17 patients who did not develop lung metastasis, only 6 (35%) showed positive staining for EFEMP1. Importantly, patients with EFEMP1-positive biopsies had a higher mortality rate of 58% compared to those with negative biopsies, whose mortality rate was 29%. Representative images of H&E and immunostaining for EFEMP1 in biopsy samples with negative, weak, moderate and strong intensities are shown in Fig. 5H. The positive staining was primarily found in the cytoplasm and membrane of OS cells.

Due to the small sample size in our study, we were unable to perform a reliable Kaplan-Meier survival analysis. Therefore, we used the recognized R2 bioinformatic platform to conduct a univariate analysis, which revealed a significant correlation between high EFEMP1 expression and poorer overall survival (246 vs. 135 months) in high-grade conventional OS patients (raw p-value=0.05, Fig 5I). We also observed a trend towards lower survival in patients with lung metastases and high EFEMP1 expression, although it did not reach statistical significance (raw p-value=0.379, Fig 5J). These findings suggest that EFEMP1 upregulation in primary tumours is associated with poor prognosis in OS patients, particularly those with high-grade conventional OS. However, further studies using larger sample sizes are required to confirm these results, as well as to understand the underlying mechanisms of EFEMP1 on cancer progression and patient outcomes.

## Discussion

In this study, we set out to understand the mechanisms by which OS cells reprogram the lung microenvironment to support the colonization and outgrowth of disseminated metastatic cells. Employing two experimental mouse models and a multi-omics approach, we demonstrated that OS cell-derived secreted factors instigate a permissive pre-metastatic lung for incoming tumour cells, comprising airways damage, ECM remodelling with fibronectin and collagen deposition by activated fibroblasts, recruitment of neutrophils and inflammation. These alterations, observed in animals carrying a primary tumour or treated with the SCR, highlight the pivotal role of OS-secreted factors in distant cell communication and in setting up the lung metastatic cascade.

Initial histological and electron microscopy examinations showed severe changes in the alveolar architecture, with septum thickening, damage in the alveolar space inflammatory infiltrates, and signs of fibrosis. The transcriptome profile revealed enrichment in multiple genes involved in related biological processes, indicating ECM remodelling and inflammation as early steps of the metastatic cascade. The IHC analysis revealed the increased deposition of fibronectin and fibrillar collagen in the premetastatic lungs of both mouse models, which was further confirmed in decellularized lung pieces. Fibrillar collagen and fibronectin form the ECM backbone and with other proteins, serve as a physical scaffold for cell adhesion and invasion by acting as ligands for numerous adhesion molecules, including the β1 and α6 integrins and CD44, which are highly expressed on 143B OS cells (Fig S6A). The adhesion assays confirmed the 143B cell attachment to these specific ECM components (Fig 3C, E), suggesting it provides the adhesion molecules for the subsequent engraftment of disseminated tumour cells.

Besides providing adhesion sites and complex signalling networks influencing cell fate, ECM remodelling is also implicated in metastatic organotropism through organ-specific integrin-ECM interactions. It has been shown that exosomal α6β4 and α6β1 integrins dictate the exosome adhesion to ECM components and lung resident fibroblasts, resulting in stromal remodelling to host metastatic cells (Hoshino *et al*, 2015). Herein, we have not explored the specific contribution of exosomes to PMN formation, but recent studies from our group confirmed that exosomes derived from 143B cells express β1 and α6 integrins and possess an intrinsic homing ability for homotypic lung metastasis (Almeida *et al*, 2022), suggesting a contribution on the lung PMN formation.

Fibroblasts are the main producers of ECM components in both homeostatic conditions and in response to injury, and the major players in the dysregulated collagen turnover and fibronectin deposition leading to tumour fibrosis or desmoplasia (DeLeon-Pennell *et al*, 2020). We showed that lung fibroblasts undergo a myofibroblast-like transition in response to tumour-secreted factors, as indicated by the increased expression of α-SMA, FAP, and vimentin with a concurrently increased deposition of fibronectin and collagen, and enhanced fibroblast proliferation. It is worth noting that fibroblasts were isolated directly from tumour-free lung tissue, without any intervening culture steps that might have impacted gene expression, thereby further validating our findings. Cancer-associated fibroblasts (CAFs) are the most prominent stromal cells and serve as positive regulators of tumour growth at the primary tumour site by reprogramming the local microenvironment (Kalluri, 2016). Similarly, metastasis-associated fibroblasts (MAFs) have been found to promote metastatic outgrowth by engaging in inflammatory-mediated communication with cancer cells (Pein *et al*, 2020; Shani *et al*, 2021). These findings were obtained in models with established lung micro-or macrometastases, while in our study the lung fibroblasts’ phenotypic changes were observed in an incredibly early stage of the metastatic cascade before tumour cell spread and colonization, which highlights the contribution of fibroblasts during the PMN formation.

The cytokine TGF-β is an inducer of myofibroblast differentiation through autocrine and paracrine mechanisms, resulting in the secretion of additional growth factors, cytokines and chemokines involved in the complex integration of signals promoting the recruitment of BMDCs, angiogenesis, immunosuppression, and ECM remodelling (Kojima *et al*, 2010; Shi *et al*, 2020). The transcriptome and RT-PCR analysis confirmed the expression of TGF-β and some other fibroblast activation inducers (e.g TNF-α and CCl2) in premetastatic lungs. These findings are in line with a study by Mazumdar *et al*. (Mazumdar *et al*, 2020b) on highly metastatic OS cell lines, including 143B, showing that OS-derived EVs were able to induce the reprogramming of lung fibroblasts through the activation of TGF-β1 and SMAD2 pathways.

Our data revealed an upregulation of the S100A8 and A9 chemoattractant in lung tissue from both mouse models, along with upregulation of TGF-β, IL-6 and TNF-α, and an increase in CD11b^+^Ly6G^+^ neutrophil infiltration and systemic expansion. These results are in line with the previous research conducted by Hiratsuka and colleagues (Hiratsuka *et al*, 2006; Hiratsuka *et al*., 2008), which demonstrated that TNF-α-induced S100A8 and A9 signalling promotes the expression of serum amyloid A3 (SAA3) and the recruitment of CD11b^+^ myeloid cells in premetastatic lungs and immunosuppression. Apparently, this inflammatory like-state accelerates the migration of primary tumour cells to the lungs. Several clinical studies have associated high levels of intratumoral neutrophils with poor prognosis in cancer patients. This correlation has been attributed to the release of pro-angiogenic and tumour-growth factors and to the suppression of immune responses (Faria *et al*, 2016; Liu *et al*, 2016; Ma *et al*, 2018). Some studies have reported the accumulation of neutrophils in future metastatic sites driven by tumour-secreted factors, but their role remains elusive as both pro-and anti-metastatic functions have been described (Charan *et al*, 2020; Granot *et al*, 2011; Kowanetz *et al*, 2010; Mazumdar *et al*., 2020a).

As part of the innate immune system, neutrophils are recruited to sites of inflammation driven by cytokines and chemotactic factors, where they participate in the inflammatory process. As already referred, the transcriptome analysis identified increased expression of several members of the CXC and CC chemokines and S100A8/A9 that are chemotactic for neutrophils, which might explain their accumulation in premetastatic lungs. Herein, we were unable to clearly identify the functional significance of neutrophils at this early step of the metastatic cascade. However, the upregulation of the immunosuppressive TGF-β in the lungs, which is the major driver of neutrophil polarization (Fridlender *et al*, 2009) supports a pro-tumorigenic phenotype, instead of providing anti-metastatic protection (Pang *et al*, 2013). The upregulation of Lcn2, a potent inducer of chemotaxis and migration of neutrophils, and downregulation of Mpo, a marker of neutrophil degranulation, from transcriptomics, support these findings.

The use of immunocompromised mice does not allow us to assess the contribution of neutrophils in the suppression of T-cell-mediated responses, which is a limitation of this study. However, inhibition of Ly6G+ cell mobilization has been shown to prevent lung metastasis formation in both immunocompetent and immunodeficient mice, suggesting a T cell-independent effect, at least in early stages (Kowanetz et al., 2010). However, further studies are needed for better clarification of the pro-metastatic contribution of neutrophils in PMN formation.

The follow-up animal studies confirmed that the reprogramming of the pulmonary microenvironment promotes and accelerates lung metastasis formation, which suggests that the intrinsic metastatic potential of tumour cells alone may not be sufficient for metastatic outgrowth and that a pre-existing growth-permissive niche is crucial for incoming cells to metastasize.

Being a systemic effect, we performed a comparative analysis of differentially secreted proteins identified in the secretome of 143B cells and the plasma samples from mice with a PT. Interestingly, the common significantly enriched GO terms were related to ECM organization, cell-matrix adhesion, neutrophil and platelet degranulation, and cytokine-mediated signalling, which are consistent with the lung microenvironmental changes previously described.

Three of the enriched proteins identified in the 143B cell secretome (EFEMP1, CRIP1 and PSMA5) were detected in the plasma of tumour mice, indicating the 143B cells as the secretory source. EFEMP1 is an ECM glycoprotein involved in organizing the ECM scaffold and promoting cell-cell and cell-ECM adhesion in normal connective tissues and disease (Timpl *et al*, 2003). This glycoprotein is involved in the carcinogenesis of some tumours, including gliomas, pancreatic, ovarian, or bladder cancer, as well as OS, by promoting cell proliferation, MMP-induced invasion, migration, EMT, and angiogenesis (Al Khader *et al*, 2022; Chen *et al*, 2021; Hu *et al*, 2009; Zhang *et al*, 2022). In a previous study, Wang *et al*. (Wang *et al*, 2015) found higher levels of EFEMP1 in the serum of OS patients compared to healthy individuals, which led them to speculate a possible link between EFEMP1 and the progression of the disease.

Our results confirmed the presence of EFEMP1 in the plasma of mice with primary OS or lung metastasis and animals treated with the 143B cell-derived SCR in a tumour-free setting, providing additional evidence for its potential role in metastatic dissemination in the initial stages of cancer onset. Although not explored, we reasoned that EFEMP1 would likely be involved in the ECM remodelling and/or participate in cell-matrix adhesion early in PMN formation, as it is part of the core effectors of these biological processes.

We analysed the expression of EFEMP1 in biopsy samples of OS patients at diagnosis to evaluate whether it could represent a risk for metastasis development. However, the small size of our patient cohort precludes us from establishing a definite causal relationship between EFEMP1 expression and the development of lung metastases. Nonetheless, we have observed that patients with EFEMP1-expressing tumours have poorer overall survival, suggesting it is associated with tumour aggressiveness, which is substantiated by the Kaplan-Meier survival analysis. Further research with larger patient cohorts are necessary to confirm this hypothesis and to gain a more comprehensive understanding of EFEMP1’s involvement in the disease.

In summary, our study sheds light on the systemic-induced lung microenvironmental changes that precede metastatic spread in OS. Integration of our data uncovers neutrophil infiltration and the functional contribution of stromal activated fibroblasts in ECM remodelling for tumour cell attachment as early pro-metastatic events, which may hold therapeutic potential in preventing or slowing the metastatic spread. Future mechanistic studies are warranted. Moreover, we identified EFEMP1 as a potential risk factor for lung metastasis and a poor prognosis factor in OS.

## Materials and Methods

### Cell culture and infection with a lentivirus encoding luciferase

The human OS cell line 143B was purchased from the American Type Culture Collection (ATCC, Manassas, USA) and cultured in Eagle’s minimum essential medium (EMEM; Sigma Aldrich, UK), supplemented with 10% heat-inactivated fetal bovine serum (FBS; Gibco, Paisley, UK), 0.015 mg/ml 5-bromo-2’- deoxyuridine (BrdU; Sigma Aldrich, St. Louis, MO, USA), 1.0 mM sodium pyruvate (Gibco; Grand Island, USA), and 1% antibiotic-antimycotic (Gibco, Grand Island, USA). The cells were maintained under standard adherent conditions in a humidified incubator with 5% CO_2_ at 37°C. Cells were authenticated by immunohistochemistry with antibodies against vimentin (Invitrogen, Thermo Fischer Scientific, Netherlands) and Ki-67 (clone MIB-1; Dako) and haematoxylin and eosin (H&E). The 143B cells were stably transduced with a lentivirus encoding Luciferase as described elsewhere (Almeida *et al*., 2022).

For secretome preparation, 143B cells (25×10^3^ cells/cm^2^) were cultured in an exosome-free culture medium for 24h. The supernatant was collected, centrifuged to remove cell debris and concentrated with Amicon Ultra-15 Centrifugal Filters (10 kDa molecular weight cut-off; Merck Millipore Ltd, Carrigtwohill, Ireland). Aliquots were stored at −80°C until usage.

### Animal studies

All animal experiments were approved by the Animal Welfare Committee of the Faculty of Medicine of the University of Coimbra and conducted following the European Community directive guidelines for the use of animals in the laboratory (2010/63/EU) transposed to the Portuguese law in 2013 (Decreto-Lei 113/2013). Athymic Swiss nude (Foxn1^nu/nu^) mice, male or female, (8-12 weeks old, 20-30 g) were purchased from the Institute for Clinical and Biomedical Research (iCBR, Coimbra, Portugal) of the Faculty of Medicine of the University of Coimbra and housed under pathogen-free conditions in individually ventilated cages, with controlled temperature/humidity (22°C/55%) environment on a 12 h light-dark cycle and with food and water ad libitum.

#### Pre-metastatic niche formation

Animals were randomized into two groups. A group was injected subcutaneously into the lower flank with 5×10^5^ of 143B-Luc^+^/100 µL PBS for primary tumour formation. The tumour growth was monitored every two days in two dimensions using a digital calliper, and mice were sacrificed when the tumour reached 50-60 mm^3^ in volume. Tumour volumes were calculated using the modified ellipsoid formula V = A × B^2^/3 (A length; B width).

The second group received a daily intraperitoneal (*i.p*.) injection of 25 μL of the 143B-derived SCR or the vehicle for 1 week, following which they were sacrificed. Animals were euthanized by cervical dislocation, and peripheral blood and lung tissue were harvested for further analysis. Peripheral blood samples were collected to K_3_EDTA tubes (Greiner bio-one, Kremsmunster, Austria) from anaesthetized animals via cardiac puncture before euthanasia. Plasma was separated by centrifugation at 3000 rpm for 10 min at 4°C, and aliquots were stored at −80°C until usage.

#### Mouse models of lung metastasis

For experimental lung metastasis formation, animals were injected intravenously (*i.v*.) into the lateral tail vein with 1.5 x 10^6^ of 143B-Luc^+^ cells/100 µL PBS. Another set of animals received a daily intraperitoneal injection of 25μL of the SCR for 1 week prior to the *i.v.* injection of 143B-Luc^+^ (1.5 x 10^6^ cells/100 uL PBS).

For spontaneous lung metastasis formation, animals were injected subcutaneously in the lower flank with 5×10^5^ of 143B-Luc^+^ cells/100 µL PBS. After reaching a maximum volume of 60-100 mm^3^, subcutaneous tumours were surgically excised and the skin incision was sutured with the animals under anaesthesia (10 mg/kg; Rompun 2%, Kiel, Germany).

Animals were monitored weekly for lung metastasis formation during a maximum of 60 days by bioluminescence imaging (BLI) on an IVIS Lumina XR (Caliper Life Sciences Inc., PerkinElmer, Massachusetts, USA). Images were acquired after *i.p.* injection of D-Luciferin (150 mg/kg), with the animals anaesthetized with 2.5% of isoflurane (Virbac, Carros, France) in 100% O_2._ Bioluminescent images were analysed using the Living Image software version 4.10 (Xenogen, Alameda, California). A region of interest (ROI) was drawn around the lesions for the quantification of the bioluminescent signal. Values are expressed photons/sec/cm^2^/sr. At the end of the study, animals were euthanized by cervical dislocation and peripheral blood and metastatic lungs were collected for histopathological analysis.

### Clinical samples

Paraffin-embedded biopsy specimens were obtained retrospectively from 29 patients diagnosed with OS. At the time diagnosis, all patients had strictly localized disease and were naïve to any chemotherapy. Of the 29 patients, 26 were classified as having conventional high-grade OS while the remaining 3 were classified as low-grade. Except for those with low-grade, all patients underwent chemotherapy prior to surgery, and all received adjuvant chemotherapy. The follow-up period ranged from 1 year to at least 10 years, or until death. The study was conducted according to the guidelines of the Declaration of Helsinki, and was approved by the Ethics Committee of the Coimbra Hospital and University Center (CHUC-021/19). Written informed consent was provided by all patients or their legal guardians.

The specific characteristics of the patients are summarized in Table 1.

**Table 1.** Clinicopathological features of osteosarcoma patients

|  |  |
| --- | --- |
| **Number of patients** | **29** |
| **Age** |  |
| Mean (range) | 33.6 (15-78) |
| **Variable** | **Number (%)** |
| **Gender** |  |
| Male | 15 (52) |
| Female | 14 (48) |
| **Primary tumour location** |  |
| Femur | 16 (55) |
| Tibia | 5 (17) |
| Humerus | 2 (7) |
| Fibula | 2 (7) |
| Others | 4 (14) |
| **Tumour grade** |  |
| High grade | 26 (90) |
| Low grade | 3 (10) |
| **Metastasis** | 12 (41) |
| **Disease-free interval (DFI)** | **Months** |
| Mean (range) | 41.4 (6-159) |

### Scanning electron microscopy (SEM)

Small fragments of resected lung tissues and decellularized matrices were fixed with 2 % glutaraldehyde and examined in a scanning electron microscope (Flex SEM 1000, HITACHI), under variable pressure scanning using an accelerating voltage of 5-10 kV.

### Transmission electron microscopy (TEM)

Resected lung tissues were sectioned into small fragments (1 mm^3^) and fixed for 2 h in 2.5% glutaraldehyde buffered with 0.1 M cacodylate buffer (pH 7.4), followed by a post-fixation in 1% osmium tetroxide (OsO_4_) for 1.5 h. After washing, samples were incubated with 1% aqueous uranyl acetate for 1 h, for contrast enhancement. Samples were then dehydrated in a graded acetone series (30-100%) followed by resin embedding using an epoxy embedding kit (Fluka Analytical, Sigma Aldrich, Germany). Ultra-thin sections were obtained with a Leica EM UC6 (Leica Co; Austria) ultramicrotome, mounted on copper grids and stained with lead citrate 0.2 % for 7 min. Images were acquired on a Tecnai G2 Spirit Bio Twin electron microscopy at 100 kV (FEI) and AnalySIS 3.2 software.

### Histopathological analysis and immunostaining

For murine models, lung tissues and primary tumours were fixed in 4% paraformaldehyde (PFA) and processed for paraffin embedding. Sections of 4 µm were stained with haematoxylin and eosin (H&E, Sigma-Aldrich, St. Louis, MO, EUA) or antibodies against fibronectin (ab2413, Abcam, USA), alpha-smooth muscle actin (α-SMA, ABT 1487, Millipore, Darmstadt, Germany) and vimentin (V9; Ventana, Arizona, EUA). Antigen retrieval was performed by immersing slides in EDTA-Tris buffer (pH 8) for 8 min at 95 °C and then blocked with a buffered hyper protein solution for 4 min to avoid nonspecific bonds. Immunostaining was performed using a Ventana Marker Platform Benchmark Ultra IHC/ISH with the resource of a multimeric indirect free biotin detection system - Optiview DAB IHC Detection Kit (Ventana Medical Systems, Arizona, EUA), according to the manufacturer instructions. A Gordon’s and Sweet silver staining was performed for the detection of reticulin fibers. Slides were observed under a light microscope Nikon Eclipse 50 I and images were captured with a Nikon-Digital Sight DS-Fi1 camera. Architectural changes were evaluated, and inter-alveolar septal thickness was measured in randomly selected H&E-stained sections.

In clinical samples, immunohistochemistry for EFEMP1/fibulin 3 (1:500, ab256457, Abcam) was performed in 4 μm tissue sections of formalin-fixed and paraffin-embedded tissue. All samples were counterstained with hematoxylin-eosin by standard methods. Immunohistochemistry positivity was considered when at least 1% of the viable neoplastic cells showed cytoplasmic and/or membrane expression of any intensity (weak, moderate or strong). The sections were assessed by an experienced pathologist.

### Primary lung fibroblasts isolation and culture

Lungs were harvested, washed in PBS, minced with scissors, and enzymatically digested with 0.1% collagenase A (Roche, Mannheim, Germany) and dispase II (2.4 U/mL; Gibco, Japan) for 90 min at 37 °C, and then filtered through a 70 µm filter strainer and washed with a physiological saline solution containing 0.05 M EDTA. The obtained cell suspension was plated into 1% gelatin pre-coated dishes and maintained in RPMI 1640 (Sigma-Aldrich, UK) supplemented with 15% FBS and 1% antibiotic-antimycotic at 37 °C under 5% CO_2_. All experiments were performed with early passages fibroblasts (P2-P4).

### Immunofluorescence

Primary fibroblasts were fixed with 4% paraformaldehyde (PFA, Sigma-Aldrich, St. Louis, MO, USA) for 20 min, permeabilized with 0.2 % TritonX-100 (Sigma-Aldrich, St. Louis, MO, USA) for 10 min, and blocked with 3% bovine serum albumin (BSA; Sigma-Aldrich, St. Louis, MO, USA) for 1 h at room temperature. Afterwards, the cells were incubated with the primary antibodies against α-SMA (1:200, ABT 1487; Millipore, Darmstadt, Germany), FAP (1:150, PA5-99313, Thermo Fisher Scientific, USA), vimentin (1:200, SP20; Thermo Fisher Scientific, USA) and fibronectin (1:200, ab2413; Abcam, Cambridge, UK) overnight at 4 °C. The fibroblasts were then incubated with secondary antibody Alexa Fluor 568 or 488 (1:200, Invitrogen, USA) for 1 h at room temperature in the dark, and nuclei were counter with 2 mg/mL Hoechst 33342 (Sigma-Aldrich, USA).

For immunofluorescence staining with Alexa Fluor 555 phalloidin, cells were fixed with PBS containing 4% sucrose and 4% paraformaldehyde for 10 min, permeabilized with 1% TritonX-100 for 10 min and blocked with 1% BSA for 45 min at room temperature, and then incubated with Alexa Fluor 555 phalloidin (1:40; Abcam, Cambridge, MA, USA) for 30 min at RT. Nuclei were stained with 2 mg/mL Hoechst 33342 (Sigma-Aldrich, Buchs, Switzerland). The chamber slides were mounted in the Vectashield Mounting Medium (Vectashield, Vector Laboratories, United States).

Decellularized lung fragments and fibroblast cell sheets were processed for immunofluorescence as described above and stained with the primary antibodies against fibronectin (1:200) and collagen IV (1:200 ab19808; Abcam, Cambridge, UK) and then incubated with secondary antibody Alexa Fluor 488 (1:200, Invitrogen, USA). Images were captured in Carl Zeiss Axio Observer Z1 inverted microscope (Carl Zeiss, Thornwood, NY) and processed using the ImageJ 1.52p software (National Institutes of Health, USA).

### Cell migration

Cell migration was analysed using the wound-healing assay. Confluent monolayers of fibroblasts were manually scratched with a 200 µL sterile pipette tip. Photographs were taken immediately after scratching (baseline) and at 6h and 24h in a Carl Zeiss Axio Observer Z1 inverted microscope (Carl Zeiss, Thornwood, NY). The wound closure at each time-point was quantified using ImageJ 1.52p software (National Institutes of Health, USA) and normalized to the baseline.

### Lung transcriptomic analysis

Total RNA was extracted from lung tissues using TRIzol reagent (Ambion by Life Technologies, USA) according to the manufacturer’s instructions. RNA was eluted in 30-40 μL RNase-free water. The concentration and quality of extracted RNA were determined with NanoDrop spectrophotometer (Thermo Scientific, Wilmington, USA). RNA was stored at 80 °C until use. Single-end sequencing was performed using the library prep kit TruSeq and the sequencing kit NovaSeq6000 SP Flowcell 100 cycles (Illumina, Inc.) for Mus musculus (Ensembl.GRCm38.82). The transcriptomic sequencing of RNA was performed on the Illumina NovaSeq 6000 instrument, at the VIB Nucleomics Core (www.nucleomics.be). The reads pre-processing was performed by VIB Nucleomics Core and involved: quality trimming (FastX 0.0.14, HannonLab), adapter trimming (cutadapt 1.15) (Martin, 2011), quality filtering (FastX 0.0.14 and ShortRead 1.40.0) (Morgan *et al*, 2009) and removal of contaminants (bowtie 2.3.3.1). The pre-processed reads were then aligned to the reference genome of Mus_musculus.Ensembl.GRCm.38.82 using STAR 2.5.2b (Dobin *et al*, 2013) and SAMtools 1.5 (Li *et al*, 2009) Bioconductor package. The expression levels of the read overlapping genes were computed using the EDASeq package (Risso *et al*, 2011) for the within-sample and between samples normalizations. The differentially expressed genes were estimated by fitting a negative binomial generalized linear model (GLM) using the edgeR 3.24.3 package (Robinson *et al*, 2010). The resultant p-value was corrected for multiple testing with Benjamini-Hochberg to control the false discovery rate (FDR).

Unless stated otherwise, the following bioinformatics analyses were conducted using the R programming language (version 4.1.1) in the RStudio integrated development environment. The volcano plots were achieved using the ggplot2 (Wickham, 2016) (version 3.3.5) and ggrepel (version 0.9.1) packages. The Venn diagrams were generated with the webtool available at bioinformatics.psb.ugent.be/webtools/Venn/. The clusterProfiler (Wu *et al*, 2021) version 4.0.5) package was used to perform the gene set enrichment analyses (GSEA) for the Gene Ontology (GO) terms and Kyoto Encyclopedia of Genes and Genomes (KEGG) pathways. Genes showing a p-value higher than 0.05 were excluded from the gene set. Heat maps were made with the ComplexHeatmap (Gu *et al*, 2016) (version 2.8.0) package. Protein-protein interaction (PPI) network of DEGs was visualized using the Search Tool for the Retrieval of Interacting Genes/Proteins (STRING) database (http://string-db.org/). Graphics were refined and figures were assembled in Adobe Illustrator 2019 (version 23.1.1).

### Flow cytometry

Fresh lung tissue was minced with fine scissors and subjected to mechanical dissociation (GentleMACS, Miltenyi Biotech, Germany). The tissue homogenates were filtered gently through a 40 μm cell strainer and centrifuged at 600g for 15 min at RT. The isolated single-cell suspension and peripheral leukocytes were fluorescently stained with anti-mouse CD11b-FITC (1:200, BioLegend, 101205) and Ly6G-PE (1:80, BioLegend, 127607). FACS Lysing solution (BD Sciences) was added to the samples for lysing red blood cells. Data were collected on a FACS Canto^TM^ II (BD Biosciences, USA) and analysed using Infinicyt V.1.8 software (Cytognos, Salamanca, Spain).

### RNA extraction and qRT-PCR

Total RNA from lung tissue was extracted using TRIzol reagent (Ambion by Life Technologies, USA) according to the manufacturer’s instructions. The concentration and quality of extracted RNA were determined using a NanoDrop spectrophotometer (Thermo Scientific, Wilmington, USA). First-strand complementary deoxyribonucleic acid (cDNA) was synthesized from 2 µg of total RNA using the NZY first-strand cDNA synthesis kit. Quantitative real-time PCR analysis was performed using a Xpert SYBR Green MasterMix (GRISP, Porto, Portugal) on a CFX Connect Real-Time PCR Detection System (Bio-Rad Laboratories, Inc). Gene expression was normalized to the housekeeping genes glyceraldehyde 3-phosphate dehydrogenase (GADPH) and hypoxanthine-guanine phosphoribosyltransferase 1 (HPRT-1) ranked as stably expressed by the RefFinder algorithm (https://www.heartcure.com.au/reffinder/). The relative expression of target genes was calculated based on the 2^−ΔΔCt^ method, where ΔΔCt=(Ct _target_−Ct _housekeeping_)_Sample_−(Ct _target_−Ct _housekeeping_)_Control_. Primer sequences (Eurofins Genomics, Lisboa, Portugal) are listed in Table 1. A Primer-BLAST search was performed to evaluate primer specificity and self-complementarity values of already validated and published primer sequences.

**Table 1:**
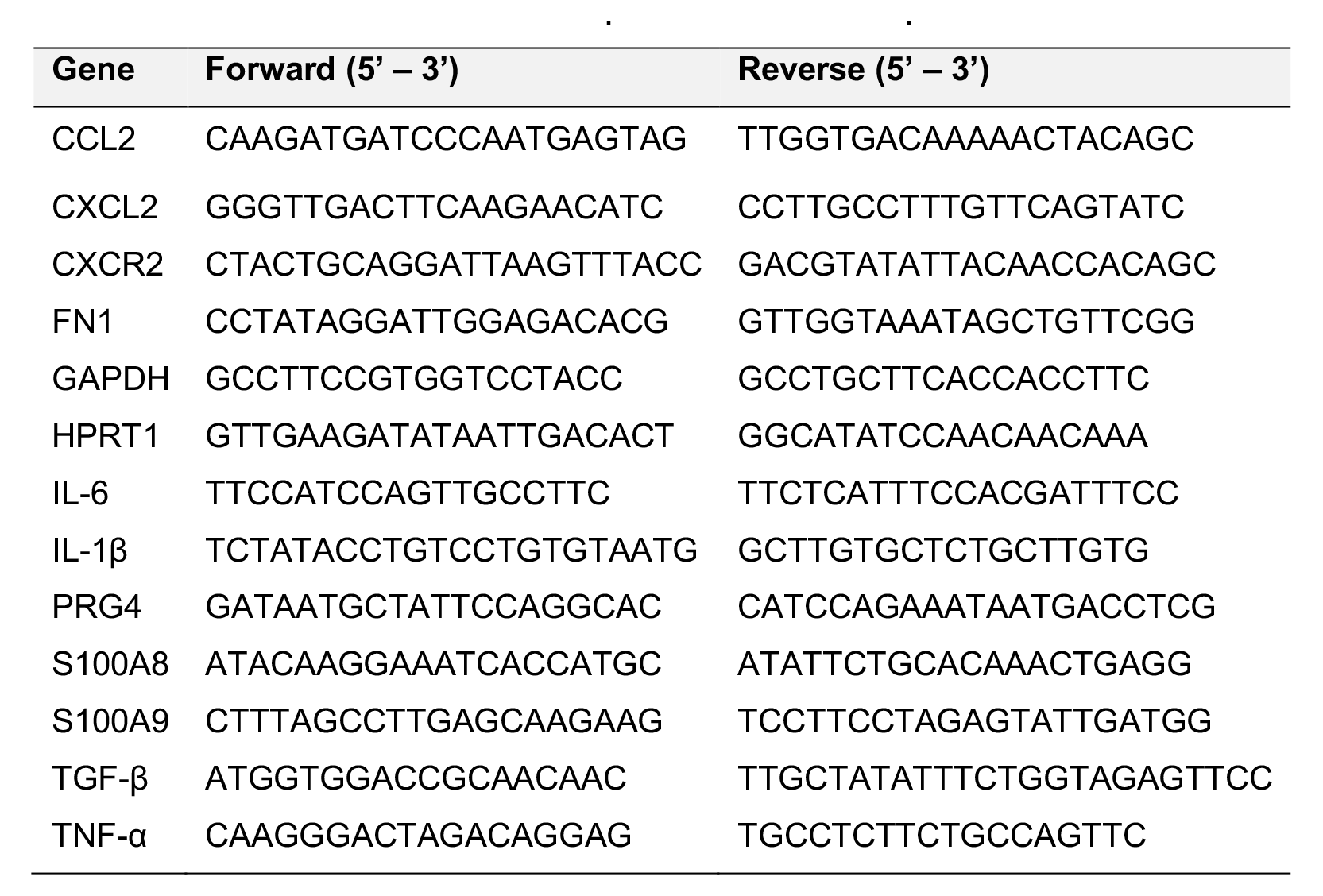
Primer sequences used in qRT-PCR

### Western Blot

Total protein extracts were prepared using standard lysis buffer, separated by sodium dodecyl sulphate/polyacrylamide gel electrophoresis (SDS-PAGE) and transferred to PVDF membranes (Bio-Rad, Hercules, CA, USA). Membranes were blocked with 5% of bovine serum albumin (BSA) in Tris-buffered saline-tween 20 (TBS-T) and were incubated with appropriate primary antibodies fibronectin (1:500; ab2413; Abcam, Cambridge, UK), collagen IV (1:500; ab19808; Abcam, Cambridge, UK), EFEMP1 (1:1000, ab256457, Abcam), ITG Β1 (1:1000, D6S1W, Cell Signalling), ITGα6 (1:1000, ab181551, Abcam) and CD44/HCAM (1:200,sc-7297, Santa Cruz Biotechnology), overnight at 4 °C. Afterward, the membranes were washed in 0.1% TBS-Tween and incubated with the secondary antibody (1:10000) for 1 h at room temperature (RT) and revealed by chemiluminescence (Clarity ECL western blotting substrates, BioRad) using ImageQuant LAS 500 chemiluminescence CCD camera (GE Healthcare Bioscience AB). Images were analysed using ImageJ software (National Institutes of Health).

### Decellularization

#### Lung fragments

Small lung fragments (2×2 mm) were incubated in hypotonic buffer (10 mM Tris HCl/0.1% EDTA, pH 7.8) for 18 h at room temperature, and then washed in PBS (3x, 1h) and immersed in a detergent solution (0.2% SDS/10 mM Tris HCl, pH 7.8) for 24 h at room temperature. Fragments were washed in hypotonic buffer (3x, 20 min) and incubated in DNAse solution (50 U/ml DNAse/ 10mMTris HCl, pH 7.8) for 3 h at 37°C. Decellularized matrices were washed in PBS (2x, 20 min) to remove the residual detergent and DNAse solution and were maintained under sterile conditions at 4°C in PBS until use. All steps were performed under agitation at 165 rpm.

#### Fibroblasts

Fibroblast cell sheets were incubated with buffer I (1M NaCl, 5 mM EDTA, 10 mM Tris/HCl pH 7.4) for 1 h at room temperature, washed with PBS (3x, 10 min) and then exposed to buffer II (0.5% [w/v] SDS, 25 mM EDTA, 10 mM Tris/HCl pH 7.4) for 30 min with agitation.

### Adhesion assays

The 96 well plates were coated with fibronectin or collagen at concentrations ranging from 1 - 10 μg/mL for 2 h. The 143B-luc^+^ cells were seeded onto the substrates and allowed to adhere for 10 min at 37°C. To remove the non-adherent cells, the wells were washed 3 times with PBS containing 1M CaCl_2_.H_2_O and MgCl_2_.6H_2_O, and then 100 μL of EMEM culture medium containing 0.3mg/mL of D-Luciferin was added to each well. The plate was read in the IVIS optical imaging system, and the bioluminescent signal was quantified using Living Image Software 4,10 (Xenogen, Alameda, CA, USA).

To evaluate the 143B cell adhesion to the decellularized lung scaffold, the decellularized fragments were placed in the 96-well plates and the 143B cells were seeded on top. The cells were allowed to adhere for 10 minutes at 37°C and the same protocol as described above was followed to quantify cell adhesion.

### Mass spectrometry

#### Protein extraction

Proteins were reduced and alkylated with 100 mM Tris pH 8.5, 1% sodium deoxycholate, 10 mM tris(2-carboxyethyl)phosphine (TCEP), and 40 mM chloroacetamide for 10 minutes at 95°C at 1000 rpm (Thermomixer, Eppendorf). 100 μg of protein were processed for proteomic analysis following the solid-phase-enhanced sample-preparation (SP3) protocol as described by Hughes *et al*. (Hughes *et al*, 2019). Enzymatic digestion was performed with Trypsin/LysC (2 μg) overnight at 37°C at 1000 rpm. The resulting peptides were cleaned-up and desalted with C18 micro columns and further quantified.

#### Proteomics data acquisition

Protein identification was performed by nanoLC-MS/MS using an Ultimate 3000 liquid chromatography system coupled to a Q-Exactive Hybrid Quadrupole-Orbitrap mass spectrometer (Thermo Scientific, Bremen, Germany). Peptides were loaded onto a trapping cartridge (Acclaim PepMap C18 100Å, 5 mm x 300 µm i.d., 160454, Thermo Scientific) in a mobile phase of 2% ACN, 0.1% FA at 10 µL/min. After 3 min loading, the trap column was switched in-line to a 50 cm by 75μm inner diameter EASY-Spray column (ES803, PepMap RSLC, C18, 2 μm, Thermo Scientific, Bremen, Germany) at 300 nL/min. Separation was achieved by mixing A: 0.1% FA, and B: 80% ACN, with the following gradient: 5 min (2.5% B to 10% B), 120 min (10% B to 30% B), 20 min (30% B to 50% B), 5 min (50% B to 99% B) and 10 min (hold 99% B). Subsequently, the column was equilibrated with 2.5% B for 17 min. Data acquisition was controlled by Xcalibur 4.0 and Tune 2.9 software (Thermo Scientific, Bremen, Germany).

The mass spectrometer was operated in data-dependent (dd) positive acquisition mode alternating between a full scan (m/z 380-1580) and subsequent HCD MS/MS of the 10 most intense peaks from the full scan (normalized collision energy of 27%). ESI spray voltage was 1.9 kV. Global settings: use lock masses best (m/z445.12003), lock mass injection Full MS, chrom. peak width (FWHM) 15s. Full scan settings: 70k resolution (m/z 200), AGC target 3e6, maximum injection time 120 ms. dd settings: minimum AGC target 8e3, intensity threshold 7.3e4, charge exclusion: unassigned, 1, 8, >8, peptide match preferred, exclude isotopes on, dynamic exclusion 45s. MS2 settings: microscans 1, resolution 35k (m/z 200), AGC target 2e5, maximum injection time 110 ms, isolation window 2.0 m/z, isolation offset 0.0 m/z, spectrum data type profile.

#### Data analysis

The raw data was processed using the Proteome Discoverer software (Thermo Scientific) and searched against the UniProt database for Homo sapiens Proteome. The Sequest HT search engine was used to identify tryptic peptides. The ion mass tolerance was 10 ppm for precursor ions and 0.02 Da for fragment ions. The maximum allowed for missing cleavage sites was set to 2. Cysteine carbamidomethylation was defined as a constant modification. Methionine oxidation and protein N-terminus acetylation were defined as variable modifications. Peptide confidence was set to high. The processing node Percolator was enabled with the following settings: maximum delta Cn 0.05; decoy database search target FDR 1%, validation based on q-value. Imputation of missing values was performed only when a peptide was detected in at least one of the replicates analyzed. Quantitative evaluation was performed by pairwise comparisons of all detected peptides and the median ratio was used for protein level comparison. Significance assessment was performed using the background-based ANOVA method implemented in Proteome Discoverer 2.2 and multiple comparison adjustment of the p-values was performed.

#### Protein functional enrichment analysis

The protein functional enrichment analyses for Gene Ontology (GO) terms and Reactome pathways were performed using the Database for Annotation, Visualization and Integrated Discovery (DAVID) 6.8 bioinformatics tool (https://david.ncifcrf.gov/summary.jsp). The data analysis and visualization of differently expressed proteins were conducted in the R programming language (version 4.1.1) in the RStudio integrated development environment. The GO and pathways plots were achieved using the ggplot2 (version 3.3.5). A basic circle packing chart with parckcircles package (available at https://r-graph-gallery.com/305-basic-circle-packing-with-one-level.html) was used for GO – biological processes. The Venn diagrams were generated with the webtool available at bioinformatics.psb.ugent.be/webtools/Venn/.

### Enzyme-linked immunosorbent assay (ELISA)

EFEMP1 ELISA was performed in mouse serum samples using Human EFEMP1 ELISA kit (Abcam, ab269552) according to the manufacturer’s instructions.

### Kaplan-Meier analysis

The Kaplan-Meier analysis was performed using the R2 database (R2: Genomics Analysis and Visualization Platform – http://r2platform.com) which contains genome-wide gene expression data of OS patient samples (dataset: Mixed OS (Mesenchymal) – Kuijjer – 127 – vst – ilmnhwg6v2). The chondroblastic, fibroblastic and osteoblastic OS patient samples (n=73) or with metastatic disease (n=37) in the database were divided into high and low EFEMP1 (ID: 2202) expressions based on scan cut-off.

### Statistical analysis

All data are expressed as mean ± standard error of the mean (SEM). Graphics and statistical analysis were performed using GraphPad Prism version 8.0.2 (GraphPad Software, San Diego, CA, USA). Independent variables were analysed by the Mann-Whitney test, whereas Kruskal-Wallis or one-way ANOVA was used for multiple comparisons. Statistical significance was set at the level of p<0.05. For the Kaplan-Meier analysis of OS patient samples, statistical differences in survival curves were calculated by log-rank (Mantel-Cox) test. Figure illustrations were created with adobe illustrator or BioRender.com.

### Data Availability

The datasets produced in this study are available in the following databases:

- RNA-Seq data: GEO database [GSE216744] (https://www.ncbi.nlm.nih.gov/geo/query/acc.cgi?acc=GSE216744)
- Protein identification: PRIDE database [PXD040814] (https://www.ebi.ac.uk/pride/archive/projects/PXD040814)

## Acknowledgements

This work was supported by the Portuguese Foundation for Science and Technology (FCT) through the project PTDC/BTM-SAL/4451/2020 and STRATEGIC PROJECTS (UIDB/04539/2020 and UIDP/04539/2020). S.A. is a PhD fellow of the FCT (PD/BDE/142929/2018). L.S. is a PhD fellow of the FCT (PD/BDE/150707/2020). G.S.-R. is a PhD fellow of the FCT (UI/BD/154407/2022). We thank T. Rodrigues from the Microscopy and Bio-Imaging Lab (iLAB) and I. Silva & S. Silva from the Flow Cytometry Core Facility.

## Author contributions

**Sara Almeida:** Conceptualization; formal analysis; investigation; methodology; writing – original draft; writing - review and editing. **Liliana Santos:** Conceptualization; investigation; methodology. **Gabriela Ribeiro:** Formal analysis; investigation; methodology. **Hugo Ferreira:** Investigation; methodology. **Nuno Lima:** Investigation; methodology. **Rui Caetano:** Investigation; formal analysis. **Mónica Abreu:** Investigation; Formal analysis. **Mónica Zuzarte:** Investigation; formal analysis. **Ana Sofia Ribeiro:** Investigation; formal analysis. **Artur Paiva:** Formal analysis. **Tânia Marques:** Investigation; methodology. **Paulo Teixeira:** Methodology**. Rui Almeida:** Investigation; Formal Analysis**. José Casanova:** Investigation**. Henrique Girão:** Investigation. **Antero J. Abrunhosa:** Investigation; supervision; funding acquisition**. Célia M. Gomes:** Conceptualization; supervision; funding acquisition; investigation; writing – review.

